# Rank- and Threat-Dependent Social Modulation of Innate Defensive Behaviors

**DOI:** 10.1101/2025.10.31.685719

**Authors:** Ling-yun Li, Xinjian Gao, Jun Zhang, Wen-wei Wu, Ya-tang Li

## Abstract

Fear and defense are among the most fundamental survival behaviors and are profoundly influenced by the social environment in group-living animals. However, it remains poorly understood how social context—and particularly dominance hierarchy, a defining feature of many social species—modulates defensive strategies under ethologically relevant conditions. To address this question, we investigated the social modulation of innate fear in mice exposed to two ethologically relevant threats: a transient visual looming stimulus and a sustained predatory threat posed by a live rat. We found that social presence alleviated threat-induced stress and modulated defensive behaviors in a rank- and threat-specific manner. During looming exposure, it reduced immediate defensive responses and alleviated post-looming anxiety, with dominants showing greater benefit. During rat exposure, it promoted a shift from passive to active defense, again most prominently in dominants. These behavioral changes were accompanied by reorganization of transitions between defensive states, indicating that social hierarchy shapes both the expression and temporal organization of innate defensive behaviors. Conversely, threat exposure strengthened social engagement, with dominant mice exhibiting more proactive social behaviors and subordinate mice responding more readily to dominant social initiations. Together, these findings demonstrate how dominance hierarchy modulates defensive responses to distinct naturalistic threats and, in turn, how threat experience shapes social behavior, providing a behavioral framework for probing the neural basis of socially modulated innate fear.

## Introduction

Innate fear of predation has been a powerful driver of the evolution of defensive strategies (Cooper and Blumstein, 2015; Lima and Dill, 1990; Mobbs et al., 2024). In group-living species, these responses are further shaped by social context (Evans et al., 2019; Krause and Ruxton, 2002). Collective behaviors—such as the circle defense of muskoxen against wolves, the coordinated retreat of meerkats to burrows, mobbing of raptors by smaller birds, and predator-specific alarm calls in primates—demonstrate how social context enhances survival by coordinating individual responses. In humans and other social animals, individuals adjust their fear perception and defensive behaviors based on conspecific cues through mechanisms such as social buffering (Kikusui et al., 2006; Qi et al., 2021), emotional contagion (de Gelder et al., 2004), social learning (Olsson and Phelps, 2007), and social appraisal (Mumenthaler and Sander, 2012). Disruptions in these processes increase vulnerability to fear-related disorders, leading to maladaptive fear responses and impaired social functioning (Grammer and Zelikowsky, 2022; Ozbay et al., 2007; Top et al., 2016). Understanding the neural mechanisms underlying the social regulation of fear is therefore crucial for treating these neurological disorders.

Most studies on the social regulation of fear have focused on learned fear paradigms (Kikusui et al., 2006; Morozov and Ito, 2019). In these paradigms, conspecific presence typically reduces fear (Davitz and Mason, 1955), yet the effect depends on partner identity and emotional state: familiar partners provide stronger buffering (Kiyokawa et al., 2014), while frightened partners can diminish or even reverse this effect (Kiyokawa et al., 2004; Morozov and Ito, 2019). While these studies have advanced our understanding of how social cues modulate fear responses, they rely on conditioning and therefore do not capture ethologically relevant threat scenarios (Cisek and Green, 2024; Dennis et al., 2021).

In contrast, innate fear paradigms provide complementary insights into naturalistic defensive re-sponses (Adolphs, 2013; Gross and Canteras, 2012; Rosen, 2004). Because innate fear responses are evolutionarily conserved, their underlying neural mechanisms are likely to generalize across individuals and species (Hein et al., 2018; Peek and Card, 2016). Recent work using such paradigms has begun to uncover how social cues regulate defensive behaviors across taxa—including flies, fish, rodents, primates, and humans—with modulation depending on the properties of threats and social cues (Blanchard and Blanchard, 1989; Coan et al., 2006; Faustino et al., 2017; Ferreira et al., 2022; Gutzeit et al., 2020; Testard et al., 2024; Vogt et al., 1981). Despite these advances, how dominance hierarchy—a fundamental organizing principle of many animal societies—regulates defensive behavior, and whether such regulation depends on the nature of the threats, remains poorly understood.

To address these questions, we developed a behavioral paradigm in which pair-housed mice with established dominance hierarchies encountered two distinct naturalistic threats: a transient visual looming stimulus that mimics the approach of an aerial predator (Yilmaz and Meister, 2013), and a sustained exposure to a live rat (Kennedy et al., 2020; Kunwar et al., 2015). Social presence alleviates threat-induced stress and modulates defensive strategies in a rank- and threat-dependent manner, attenuating looming-evoked defense while promoting active coping during rat exposure, with the effects more pronounced in dominants. Furthermore, shared threat experience reinforces social roles and strengthens social engagement. Together, these findings reveal reciprocal interactions between defensive and social behaviors across distinct threat contexts, providing a behavioral framework for future investigations into the neural mechanisms underlying social modulation of innate fear.

## Results

### Social presence attenuates looming-evoked defensive behavior in a rank-dependent manner

Building on our recent findings that dominant and subordinate mice adopt distinct defensive decision-making strategies in response to looming threats (Li et al., 2025), here we investigate how these decisions are shaped by social context within an established dominance hierarchy. We developed a behavioral paradigm in which mice were exposed to looming stimuli either alone or with a social partner (Figure 1A). Specifically, adult male sibling mice were pair-housed for two weeks to establish stable dominance hierarchies, which were assessed using the tube test (Fan et al., 2019). Following habituation to the arena, each mouse was tested in both single and paired conditions. Dominance status was reassessed after threat exposure; only one pair showed a rank reversal and was excluded from subsequent analyses.

**Figure 1:**
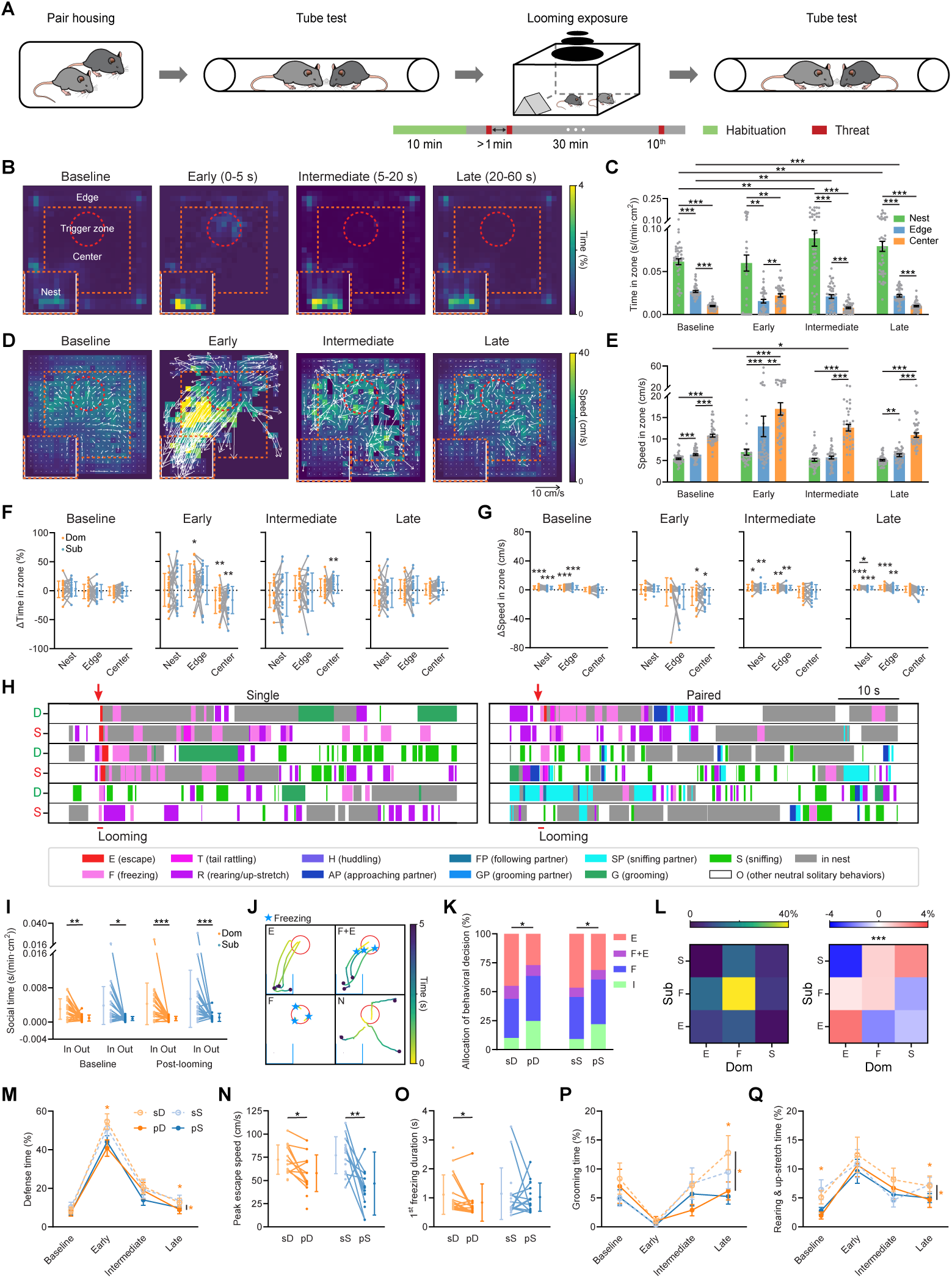
Social presence attenuates looming-evoked defensive behavior in a rank-dependent manner. (A) Schematic of the looming exposure paradigm. (B) Heatmaps of average time allocation in nest, edge, and center zones during different phases under single exposure, *n* = 40 mice. (C) Time spent in each zone per minute, normalized by zone area. *n* = 40 mice, paired *t*-test. (D) Heatmaps of average locomotion speed under single exposure. Arrows indicate mean velocity vectors, *n* = 40 mice. (E) Locomotion speed in each zone. *n* = 40 mice, paired *t*-test. (F) Change in zone occupancy (paired - single). *n* = 20 mice for both dominant (Dom) and subordinate (Sub) groups, paired *t*-test. (G) Change in locomotion speed (paired - single). *n* = 20 mice for both groups, paired *t*-test. (H) Representative raster plots of behaviors during single and paired looming exposures. D: dominant; S: subordinate. (I) Distribution of social time inside and outside of the nest. *n* = 20 mice for both groups, paired *t*-test. (J) Representative trajectories showing distinct behavioral decisions within 5 s after looming onset. (K) Distribution of behavioral decisions following looming exposure. *n* = 89, 85, 88, 96 trials for sD (single dominant), pD (pair dominant), sS (single subordinate), and pS (pair subordinate) groups, respectively; chi-squared test. (L) Left, expected behavior co-occurrence probability. Right, observed minus expected co-occurrence probability assuming independence. *n* = 20 mice for both groups. (M) Time spent in defensive behaviors (freezing, escape, tail-rattling, and rearing/up-stretch) during different phases. *n* = 20 mice for both groups, two-way ANOVA with post hoc Tukey’s range test. (N) Peak escape speed. *n* = 16, 18, 15, 15 mice, paired *t*-test. (O) Duration of the first looming-evoked freezing bout. *n* = 16, 19, 15, 19 mice, paired *t*-test. (P) Time spent on self-grooming. *n* = 20 mice per group, two-way ANOVA with post hoc Tukey’s range test. (Q) Time spent on rearing and up-stretch. *n* = 20 mice per group, two-way ANOVA with post hoc Tukey’s range test. \**p* < 0.05; ** *p* < 0.01; *** *p* < 0.001.

During baseline, mice showed a strong preference for the nest and, when exploring the arena, stayed close to the arena walls, consistent with thigmotaxis in the open-field test (Figures 1B–C and S1A–B). Looming stimuli were triggered when mice entered a defined trigger zone in the center of the arena. While looming stimuli elicited immediate defensive responses, their impact on behavior and emotional state persists beyond the initial reaction. To capture both the immediate and sustained effects, we analyzed behavior across three temporal windows relative to looming onset: an early phase (0–5 s), an intermediate phase (5–20 s), and a late phase (20–60 s). The early phase was defined to capture the initial defensive responses, as looming-evoked freezing typically lasts less than 5 s (De Franceschi et al., 2016; Li et al., 2025; Yang et al., 2020; Yilmaz and Meister, 2013), and the intermediate and late phases were defined to distinguish the acute-to-sustained transition from sustained behavior.

Looming exposure altered both spatial occupancy and locomotion speed, with distinct effects across the three temporal phases (Figures 1B–E). In the early phase, the choice between freezing and escape produced bimodal distributions of both nest occupancy (*p <* 0.001, Kolmogorov–Smirnov (KS) test) and center-zone speed (*p <* 0.001, KS test). In the intermediate phase, many mice that initially froze subsequently escaped to the nest and entered a prolonged freezing state, resulting in increased locomotion speed in the center zone. In the late phase, behavior shifted toward a sustained anxiety-like state characterized by increased nest occupancy.

The presence of a social partner during the baseline did not change spatial occupancy, but promoted locomotion in the nest and edge zones (Figures 1F–G), suggesting that social presence attenuates anxiety-related suppression of movement, an effect that persists into the post-looming period. The lack of change in spatial occupancy, in contrast, may reflect competing influences: reduced anxiety promotes exploration outside the nest, whereas social opportunities motivate mice to remain inside.

To test this hypothesis and further assess how social context modulates defensive behavior, we manually annotated behaviors (Figures 1H and S1C). As predicted, mice spent more time engaged in social interactions within the nest (Figure 1I). Furthermore, mice exhibited fewer defensive behaviors under paired conditions. To quantify this social modulation, we classified defensive responses during the early phase into four decision types: escape (E), escape after assessment (F+E), freezing (F), and no response (N) (Figure 1J, Videos 1–4). Social presence significantly altered decision patterns in both dominant and subordinate mice (Figure 1K). Interestingly, freezing, escape, and social behaviors were temporally correlated between dominant and subordinate mice (Figure 1L). Because looming stimuli were rarely triggered by the co-presence of both mice in the trigger zone (Figure S1D), this correlation cannot be attributed to concurrent threat exposure, but instead suggests that social partners dynamically influence each other’s defensive responses.

Notably, the modulatory effects depended on social rank. Total defensive time decreased significantly only in dominant mice (Figure 1M). Rank-dependent modulation was also observed in escape speed and freezing duration: subordinates showed a greater reduction in escape speed (Figure 1N), while only dominants showed a shortened freezing duration (Figure 1O). Rank-dependent modulation persisted into the late phase. Under social conditions, stress-induced self-grooming was alleviated predominantly in dominant mice (Figures 1P and S1E). A similar rank-dependent reduction was observed for anxiety-associated rearing and up-stretch behaviors (Figure 1Q).

Together, these results demonstrate that social context shapes both immediate and sustained defensive responses to transient visual threats in a rank-dependent manner, modulating not only behavioral decisions but also the expression of specific defensive and anxiety-like behaviors.

### Social presence alleviates sustained predatory stress and fine-tunes defensive strategies across ranks

In natural environments, prey must survive both sudden, unpredictable predatory strikes and the sustained presence of predators; the latter imposes prolonged stress and elicits a broader repertoire of defensive behaviors. To determine whether the effects of social context generalize across distinct forms of threat, we next examined defensive responses during live-rat exposure (Kennedy et al., 2020)(Figure 2A). Following a 5-minute habituation, a live rat was placed in an adjacent perforated, transparent chamber, allowing multisensory predatory cues to reach the mouse for 5 minutes.

**Figure 2:**
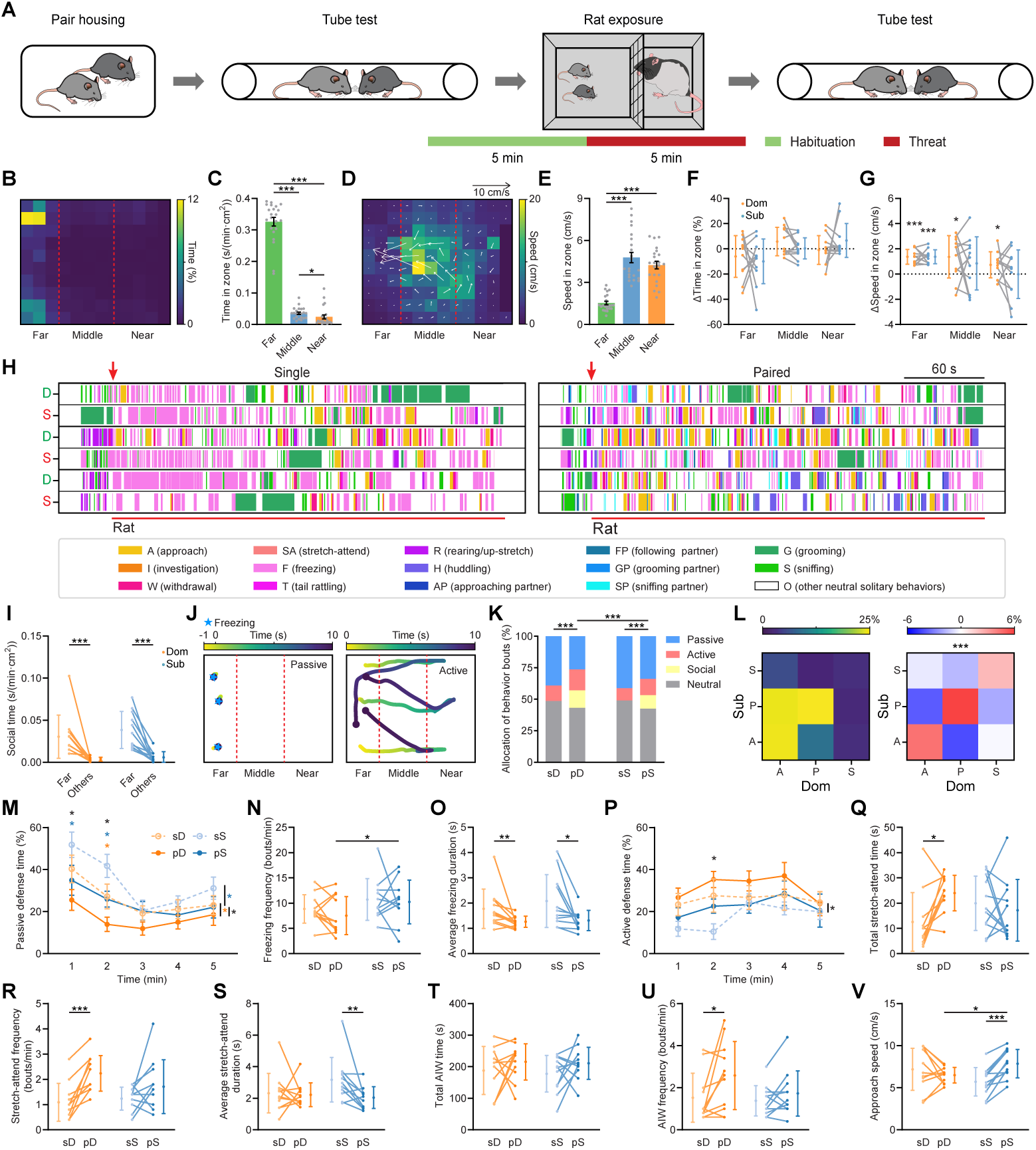
Social presence alleviates sustained predatory stress and fine-tunes defensive strategies across ranks. (A) Schematic of the rat exposure paradigm. (B) Heatmap of average time allocation in far, middle, and near zones during rat exposure, *n* = 22 mice. (C) Time spent in each zone per minute, normalized by zone area. *n* = 22 mice, paired *t*-test. (D) Heatmap of average locomotion speed, with arrows indicating average velocity vectors, *n* = 22 mice. (E) Speed in each zone. *n* = 22 mice, paired *t*-test. (F) Difference (paired - single) in time spent across zones during rat exposure. *n* = 11 mice for both dominant and subordinate groups, paired *t*-test. (G) Difference (paired - single) in speed across zones during rat exposure. *n* = 11 mice for each group, paired *t*-test. (H) Representative raster plots of behaviors during single and paired live rat exposure. D: dominant; S: subordinate. (I) Distribution of social time in far and other zones. *n* = 11 mice for both dominant and subordinate groups. (J) Representative trajectories of example mice displaying passive (left) and active (right) defense during rat exposure. (K) Distribution of behavioral categories during rat exposure. *n* = 11 mice for each group, chi-squared test. (L) Left: expected behavior co-occurrence probability. Right: observed minus expected co-occurrence probability assuming independence. *n* = 11 mice for each group. (M) Time spent on passive defense across time windows. *n* = 11 mice for each group, two-way ANOVA with post hoc Tukey’s range test. (N) Freezing frequency. *n* = 11 mice for each group, paired *t*-test. (O) Average freezing duration. *n* = 11 mice for each group, paired *t*-test. (P) Time spent on active defense across time windows. *n* = 11 mice for each group, two-way ANOVA with post hoc Tukey’s range test. (Q–S) Total stretch-attend time (Q), frequency (R), and average duration (S). *n* = 11 mice for each group, paired *t*-test. (T–V) Total AIW (approach-investigation-withdrawal) time (T), frequency (U), and approach speed (V). *n* = 11 mice for each group, paired *t*-test. \**p* < 0.05; ** *p* < 0.01; *** *p* < 0.001.

Mice differentially allocated their time across spatial zones relative to the predator (Figures 2B–C, S2A–C), and locomotion speed peaked in the middle zone when animals moved toward the far zone (Figures 2D–E). These patterns suggest a gradient of perceived risk: highest near the predator, moderate in the middle zone, and lowest in the far zone. Echoing our looming exposure findings, a social partner increased locomotion speed in the far zone (Figure 2G), reflecting reduced anxiety, but did not change time allocation across zones (Figure 2F), consistent with competing influences between exploration and social interaction (Figure 2I).

Manual behavioral annotation revealed four new active defensive behaviors during rat exposure: stretch-attend—an active risk-assessment posture—and a fixed behavioral motif comprising approach, investigation, and withdrawal (AIW) (Figures 2H and S2D). Freezing and tail rattling were grouped as passive defensive behaviors, and together with social behaviors and neutral solitary behaviors, formed four behavioral categories (see Methods) (Figures 2J–K, Videos 5–8). Social presence altered the expression of individual behaviors (Figure 2H) and reshaped the distribution across these categories in both ranks, amplifying rank-dependent differences (Figure 2K). Furthermore, consistent with the results under the looming paradigm, active defense, passive defense, and social behaviors were temporally correlated between dominant and subordinate mice (Figure 2L), supporting behavioral influences between social partners.

The amplification of rank difference from single to paired conditions was driven by a greater shift from passive to active defense in dominant mice. Social presence reduced passive defense time in both ranks and generated a rank difference under paired conditions (Figure 2M). This difference was reflected in freezing frequency and a greater reduction in freezing duration in dominant mice (Figures 2N–O and S2E–F). Social presence also produced a rank difference in active defense time under paired conditions (Figure 2P). Specifically, it increased the total time spent in stretch-attend behavior and its frequency mainly in dominant mice (Figures 2Q–R), while reducing the average duration of stretch-attend bouts in subordinate mice (Figure 2S). For the AIW motif, social presence selectively increased its frequency in dominant mice without altering its total time or average duration (Figures 2T–U and S2G). In contrast, it increased approach speed, but not withdrawal speed, in subordinate mice (Figure 2V and S2H).

These results indicate that social presence alleviates sustained predator-induced stress and reshapes defensive strategies in a rank-dependent manner. Dominant mice show a broad shift from passive to active defense across behavioral measures, whereas subordinate mice exhibit more selective modulation of active defense. These effects parallel those observed under looming stimulation, suggesting that social modulation of defensive behaviors generalizes across threat types.

Rank- and threat-dependent social modulation of defensive behavioral transitions Having established that social context reduces defensive behavior in a rank- and threat-dependent manner, we next asked how social context influences transitions between distinct behaviors. To address this, we constructed behavioral transition maps for both social ranks. These maps visualize the sequential structure of behaviors and illustrate how social context reconfigures behavioral sequences across the dominance hierarchy under distinct threat conditions (Figure 3).

**Figure 3:**
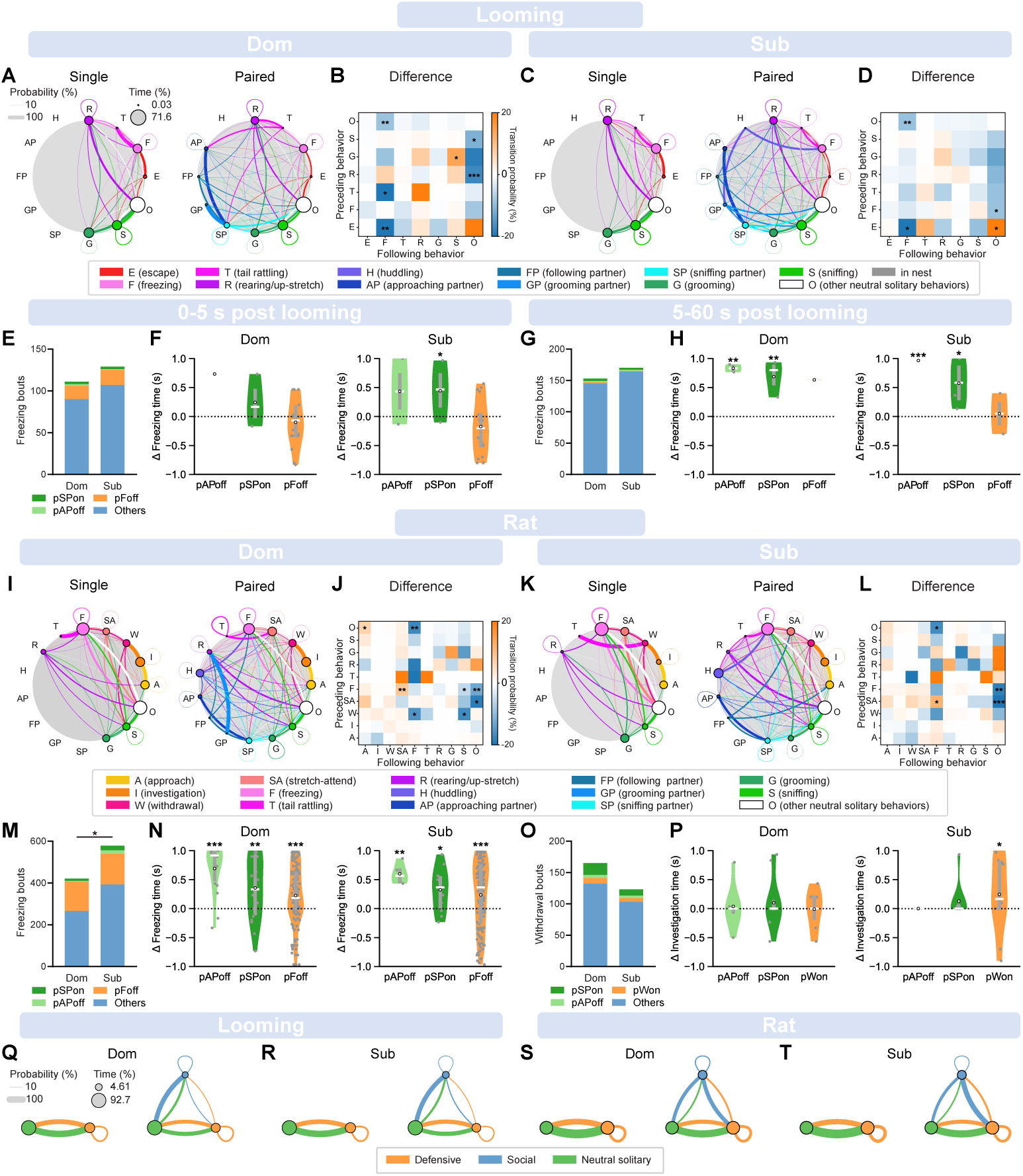
Rank- and threat-dependent organization of defensive behavioral transitions. (A) Behavioral transition maps during single (left) and paired (right) looming exposure in dominant mice. Node size scales with total time allocated to each behavior; line thickness scales with transition probability. *n* = 20 mice, paired *t*-test. (B) Matrix of behavioral transition differences between single and paired conditions (paired *−* single). *n* = 20 mice, paired *t*-test. (C–D) Same as (A–B) for subordinate mice. *n* = 20 mice, paired *t*-test. (E) Distribution of freezing bouts during looming exposure. Colors indicate freezing bouts that overlapped with the partner’s sniffing-partner onset (pSPon, dark green), approaching-partner offset (pAPoff, light green), freezing offset (pFoff, orange), and all other behaviors (Others, blue). *n* = 111 bouts (Dom) and 129 bouts (Sub). (F) Change in freezing time (pre *−* post) within a 1-s window aligned to partner’s behavior during the 0–5 s post-looming period in dominant (left) and subordinate (right) mice. Dominant: *n* = 1 (pAPoff), 3 (pSPon), and 18 (pFoff); subordinate: *n* = 2 (pAPoff), 3 (pSPon), and 16 (pFoff), permutation test. (G) Same as (E) for the 5–60 s post-looming period. *n* = 153 bouts (Dom) and 170 bouts (Sub). (H) Same as (F) for the 5–60 s post-looming period. Dominant: *n* = 2 (pAPoff), 3 (pSPon), and 1 (pFoff); subordinate: *n* = 2 (pAPoff), 4 (pSPon), and 2 (pFoff), permutation test. (I–L) Same as (A–D) for rat exposure. *n* = 11 mice per rank, paired *t*-test. (M) Distribution of freezing bouts during rat exposure. Color codes as in (E). *n* = 421 bouts (Dom) and 578 bouts (Sub). (N) Change in freezing time (pre *−* post) within a 1-s window aligned to partner’s behavior during rat exposure in dominant (left) and subordinate (right) mice. Dominant: *n* = 18 (pAPoff), 22 (pSPon), and 164 (pFoff); subordinate: *n* = 5 (pAPoff), 11 (pSPon), and 153 (pFoff), permutation test. (O) Distribution of withdrawal bouts during rat exposure. Colors indicate withdrawal bouts that overlapped with partner’s sniffing-partner onset (pSPon, dark green), approaching-partner offset (pAPoff, light green), withdrawal onset (pWon, orange), and all other behaviors (Others, blue). *n* = 165 bouts (Dom) and 123 bouts (Sub). (P) Change in investigation time (pre *−* post) within a 1-s window aligned to partner’s behavior during rat exposure in dominant (left) and subordinate (right) mice. Dominant: *n* = 7 (pAPoff), 11 (pSPon), and 7 (pWon); subordinate: *n* = 5 (pAPoff), 8 (pSPon), and 13 (pWon), permutation test. (Q) Transition maps of behavioral categories for single (left) and paired (right) looming exposure in dominant mice. *n* = 20 mice. (R) Same as (Q) for subordinate mice. *n* = 20 mice. (S–T) Same as (Q–R) for rat exposure. *n* = 11 mice per rank. \**p* < 0.05; \*\**p* < 0.01; *** *p* < 0.001.

Three key patterns emerged. First, defensive strategies exhibited clear threat-specific transition structures under single-exposure conditions. During looming exposure, escape-to-freezing transitions were frequent in both dominants (59.3%) and subordinates (59.4%), and freezing-to-rearing transitions occurred in approximately one-fourth of freezing bouts (25.6% in dominants and 21.9% in subordinates) (Figures 3A, 3C, S3A, S3C). This sequential progression suggests heightened vigilance following defense against a transient threat. In contrast, during rat exposure, withdrawal-to-freezing transitions were often observed (29.6% in dominants and 36.3% in subordinates), and about one-third of these bouts transitioned to subsequent freezing (freezing-to-freezing; 24.1% in dominants and 34% in subordinates), indicative of a sustained defensive state under persistent predatory stress (Figures 3I, 3K, S3E, S3G). Furthermore, transitions during rat exposure confirmed the AIW motif (dominants: 95.6%, 93.8%; subordinates: 89.2%, 95.5%), highlighting an active threat-evaluation strategy that was absent during looming exposure.

Second, social presence modulated transitions among defensive behaviors in a threat- and rank-dependent manner. During looming exposure, social presence reduced escape-to-freezing transitions in both ranks, consistent with reduced fear levels (Figures 3A–D, S3A–D). During rat exposure, withdrawal-to-freezing transitions were reduced in both ranks, but reached significance only in dominant mice. Consistently, social presence increased transitions from freezing to stretch-attend—an active risk-assessment posture—again selectively in dominant animals (Figures 3I–L, S3E–H). This pattern aligns with the passive-to-active shift observed predominantly in dominants (Figures 2M, 2P). Notably, social presence did not affect the approach–investigation–withdrawal motif, indicating that this sequence represents a stable behavioral module.

Third, social presence modulates the termination of defensive behaviors dependent on both threat level and social rank. During looming exposure, about 10% of freezing bouts transitioned to approaching or sniffing a partner in both ranks (Figures S3B and S3D), suggesting that social interactions can facilitate freezing termination. In addition, partner freezing offset may also contribute to freezing termination of the focal mouse (Figure 1L). To quantify the partner’s influence, we identified freezing bouts during which the partner stopped approaching the focal mouse, began sniffing the focal mouse, or terminated its own freezing. In both ranks, freezing duration was significantly reduced, with stronger effects during the late phase than the early phase (Figures 3E–H, S3I–J). During rat exposure, about 10% freezing was followed by huddling (Dom: 8.3%, Sub: 15.7%, Figures S3F and S3H); conversely, about half huddling bouts were followed by freezing (Dom: 39.4%, Sub: 52.1%), suggesting social interactions interleaved with, rather than terminated, ongoing defensive states. Consistently, partner behaviors significantly shortened freezing duration in about one third of freezing bouts in both ranks (Figures 3M–N, S3K). For termination of the AIW active defense, defined by withdrawal onset, only subordinate mice were influenced by the partner’s withdrawal (Figures 3O–P, S3L).

This threat-specific embedding of social behaviors was also evident at the category level: during rat exposure, social behaviors were more likely to transition into defensive behaviors than during looming exposure (Figures 3Q–T), consistent with a more sustained fear state. These results indicate that threat type not only shapes defensive strategies but also reorganizes how social interactions are integrated into ongoing behavioral sequences in a rank-dependent manner.

### Threat exposure reinforces social roles and promotes cohesive behavior

Having established how social context shapes defensive responses to distinct threats, we next asked the reciprocal question: how does threat exposure influence social behavior? Looming exposure selec-tively increased total social time and average social bout duration in subordinate mice (Figures 4A–C). In contrast, rat exposure increased both measures across ranks while reducing the frequency of social interactions (Figures 4D–F), indicating a shift toward prolonged social engagement under sustained threats. Consistently, huddling emerged as the predominant social behavior during rat exposure in both ranks (Figures 4G, S4A–C), likely reflecting a shift toward coordinated proximity that reduces individual risk and enhances collective security. Both looming and rat exposure also suppressed exploratory behaviors, an effect largely alleviated by the presence of a social partner (Figures S4H–M).

**Figure 4:**
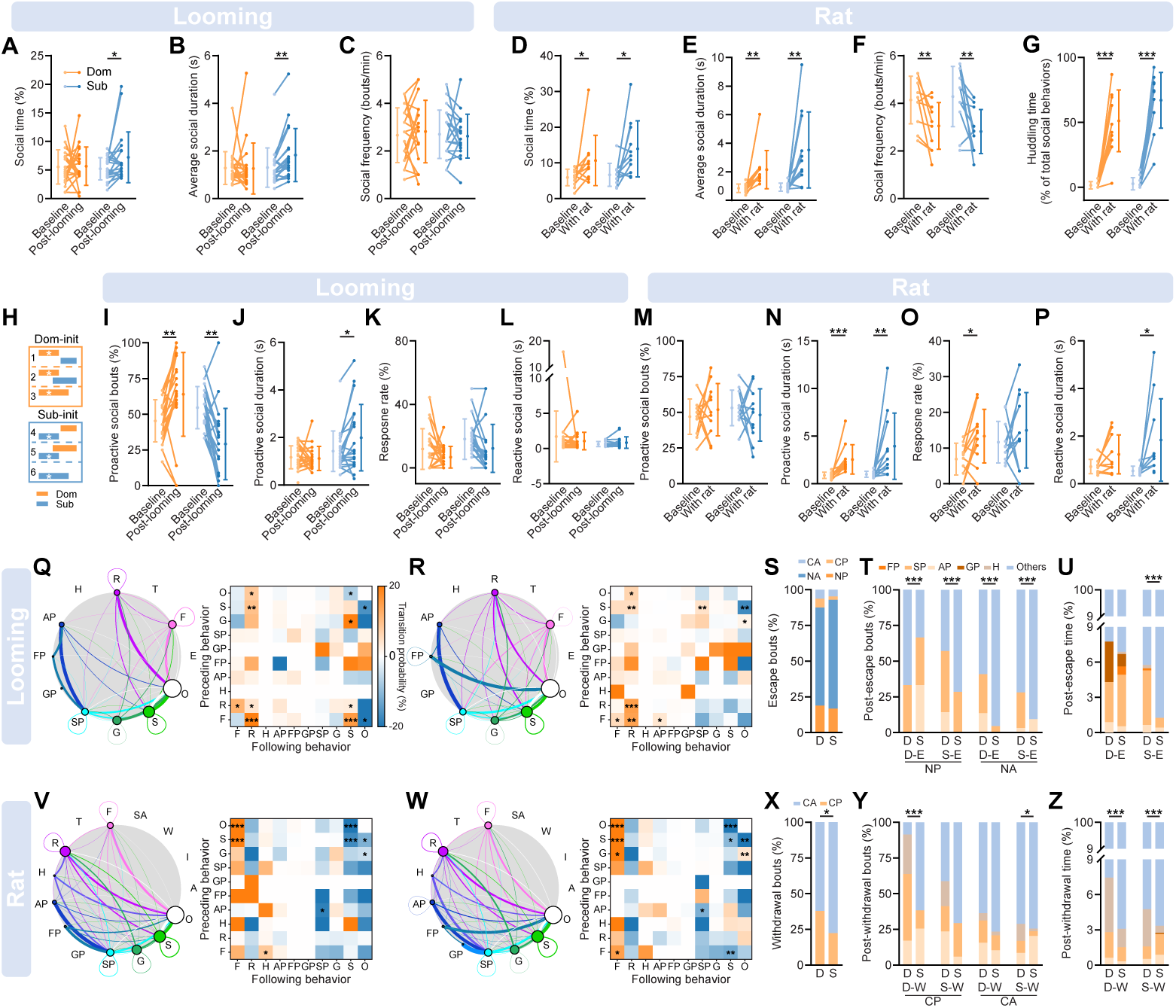
Threat exposure reinforces social roles and promotes cohesive behavior. (A) Time spent on social behaviors within a 1-minute window before and after looming exposure. *n* = 20 mice for both groups, paired *t*-test. (B) Average bout duration of social behaviors before and after looming exposure. *n* = 20 mice for both groups, paired *t*-test. (C) Frequency of social behaviors before and after looming exposure. *n* = 20 mice for both groups. (D) Time spent on social behaviors during a 5-minute window before and during rat exposure. *n* = 11 mice for both groups, paired *t*-test. (E) Average bout duration of social behaviors before and during rat exposure. *n* = 11 mice for both groups, paired *t*-test. (F) Frequency of social behaviors before and during rat exposure. *n* = 11 mice for both groups, paired *t*-test. (G) Huddling time expressed as a percentage of total social behavior time during a 5-minute window before and during rat exposure. *n* = 11 mice for both groups, paired *t*-test. (H) Schematic illustration of social interactions. Asterisks (*) denote proactive social behaviors. Bouts 1, 2, 4, and 5 represent proactive bouts that elicited a response from the partner, whereas bouts 3 and 6 represent the ones that failed to elicit a partner’s response. (I–L) Social interactions in the looming paradigm: (I) Percentage of proactive social bouts across ranks. (J) Average duration of proactive bouts. (K) Response rate to proactive bouts. (L) Average duration of reactive bouts. *n* = 20 mice for each group, paired *t*-test. (M–P) Same plots as I–L for the rat paradigm. *n* = 11 mice for each group. (Q) Left: behavior transition maps during baseline for paired looming exposure for dominant mice. Right: Matrix of behavior transition differences between threat and baseline conditions (threat *−* baseline). *n* = 20 mice, paired *t*-test. (R) Same plot for subordinate mice in looming exposure. *n* = 20 mice, paired *t*-test. (S) Escape bouts categorized by termination status: CA, corner alone; CP, corner paired; NA, nest alone; NP, nest paired. “Alone” indicates the partner was absent from the destination location at bout termination, whereas “paired” indicates the partner was present. *n*= 32 bouts (Dom) and 42 bouts (Sub). (T) Distribution of the first behavior following escape. D-E and S-E denote escape bouts performed by dominant and subordinate mice, respectively. *n* = 6, 7, 22, 32 bouts, Fisher’s exact test. (U) Time allocation of different behaviors during a 10-s time window following looming exposure. *n* = 32 bouts (D-E) and 42 bouts (S-E), Fisher’s exact test. (V–W) Same plot as Q–R for rat exposure. *n* = 11 mice, paired *t*-test. (X–Z) Same plot as S–U for withdrawal bouts in rat exposure. (X) *n*= 124 bouts (Dom), 76 bouts (Sub), chi-squared test. (Y) *n* = 47, 17, 77, 59 bouts, Fisher’s exact test. (Z) *n* = 160 bouts (D-W) and 120 bouts (S-W), Fisher’s exact test. \**p* < 0.05; ** *p* < 0.01; *** *p* < 0.001.

To determine how threat exposure alters the dynamics of social interaction, we classified social bouts as either proactive or reactive (Figure 4H, see Materials and Methods). Looming exposure increased proactive social bouts in dominant mice but decreased them in subordinates (Figure 4I), whereas rat exposure produced no rank difference (Figure 4M). Threat exposure also prolonged proactive social bouts in subordinates after looming exposure and in both ranks during rat exposure (Figures 4J and 4N). During rat exposure, proactive social initiations by dominant mice were more likely to elicit responses from subordinate mice (Figures 4K and 4O), which in turn exhibited longer reactive social bouts (Figures 4L and 4P). Because huddling accounted for about half of total social time, we performed the same analysis for huddling behavior. Subordinate mice exhibited longer proactive huddling bouts than dominant mice (Figures S4D–G), suggesting a greater tendency to seek affiliative contact under sustained threat.

Next, we ask how threat exposure reshapes the structure of behavioral transitions. After looming exposure, freezing-to-rearing and rearing-to-rearing transitions increased significantly in both ranks (Figures 4Q–R), likely reflecting heightened vigilance after exposure to a transient threat. We then specifically examined how mice transitioned from escape to subsequent social interactions. After escape, dominant mice tended to flee toward their partner more often than subordinates, regardless of whether the partner was in the nest or the corner (Figures 4S and S4N). When escape ended with both mice together in the nest, the first post-escape social interaction was more often initiated by the partner than by the escaping mouse (Figure 4T, NP). In contrast, when the escaping mouse reached the nest alone, dominant mice were more likely than subordinate mice to initiate social behavior, regardless of which animal had escaped (Figure 4T, NA). Thus, post-escape social interactions depended on the spatial relationship between the two mice: partners preferentially initiated social contact when already together, whereas dominant mice actively re-established social contact when separated. Consistent with this pattern, escape by dominant mice increased social behavior during the subsequent 10 s in both ranks, whereas following subordinate escape, dominant mice displayed more social behavior than subordinate mice (Figures 4U and S4P–Q).

Under rat exposure, huddling-to-freezing transitions increased in both ranks (Figures 3I and 3K), and freezing-to-huddling transitions increased in dominant mice (Figures 4V–W), indicating more frequent switching between defensive and social states during sustained threat. We next examined the social interactions immediately following withdrawal from the rat. After withdrawal, partners were more likely to remain in the far zone (Figure S4O), and dominant mice were more likely than subordinate mice to withdraw toward their partner (Figure 4X). Across all withdrawal bouts, dominant mice were consistently more likely to initiate social behavior following withdrawal by either animal (Figure 4Y), indicating preferential recruitment of subordinate partners after predator investigation. Consistent with this interpretation, dominant mice displayed more social behavior than subordinate mice during the subsequent 10 s under all four withdrawal conditions (Figures 4Z and S4R–S).

Overall, these findings demonstrate that naturalistic threats reshape social behavior in a threat-and rank-dependent manner. Whereas transient looming selectively increased social engagement in subordinate mice, sustained predator exposure promoted prolonged affiliative interactions, particularly huddling, across both ranks. Across both paradigms, dominant mice consistently initiated post-threat social interactions and recruited subordinate partners, suggesting that dominant individuals play a central role in coordinating social engagement following threat exposure.

## Discussion

Our study establishes a behavioral framework for investigating how dominance hierarchy shapes innate defensive behaviors under distinct naturalistic threats. Using transient looming stimuli and sustained rat exposure to simulate distinct predator encounters, we found that the presence of a social partner modulates threat-induced behaviors in a rank- and threat-dependent manner. During looming exposure, social presence reduced acute defensive responses and promoted faster recovery, with dominants showing stronger modulation (Figure 1). During rat exposure, social presence promoted a shift from passive to active defense, again more prominently in dominants (Figure 2). These effects were accompanied by changes in the structure of behavioral transitions (Figure 3), suggesting that social hierarchy influences not only the expression of defensive behaviors but also their temporal organization. Threat exposure, in turn, reshaped social interactions, with dominant mice exhibiting more proactive social behaviors and subordinate mice becoming increasingly responsive to dominant initiations (Figure 4).

These findings broaden our understanding of how social context regulates fear responses, which has largely focused on learned fear paradigms (Kikusui et al., 2006; Morozov and Ito, 2019). Our finding that conspecific presence reduces innate fear is consistent with evidence across species, including zebrafish (Faustino et al., 2017), flies (Ferreira et al., 2022), rats (Bowen et al., 2013), monkeys (Testard et al., 2024), and humans (Coan et al., 2006). Nevertheless, how social hierarchy modulates innate fear has remained poorly understood. One previous study in rats reported that dominance status influences animal behavior during predator exposure (Blanchard and Blanchard, 1989), potentially reflecting increased stress in subordinate animals (Blanchard and Blanchard, 1990; Blanchard et al., 1993). Yet prior work lacked paired versus single comparisons, leaving the role of social context unresolved. Our work fills this gap and demonstrates that defensive responses are jointly shaped by threat type and social rank, emphasizing survival-critical behaviors beyond fear expression alone.

An important issue is whether rank- and threat-dependent differences arise from differences in perceived danger or from distinct behavioral strategies. In complex natural environments, animals have evolved a rich repertoire of defensive strategies to maximize survival under varying ecological challenges (Mobbs et al., 2024). Even when faced with the same threat, defensive decisions are shaped by prior experience, internal state, social context, and other contextual factors (Evans et al., 2019; Li et al., 2025). We propose that the observed behavioral differences arise from rank-dependent modulation of threat perception through social buffering and threat-dependent selection of defensive strategies.

Across both paradigms, social presence consistently reduced threat-induced stress and reshaped defensive behavior, with dominant mice showing stronger modulation. However, the behavioral consequences of social modulation differed depending on the threat type. Under looming exposure, which represents a transient and unpredictable visual threat, social presence primarily suppressed immediate defensive responses and alleviated post-threat anxiety, thereby facilitating behavioral recovery. Consistent with reduced fear and faster recovery, social presence decreased escape-to-freezing transitions (Figures 3A–D). In contrast, under rat exposure, which represents a sustained and predictable multisensory threat, social presence promoted a shift from passive to active defensive strategies rather than simply suppressing defensive responses. Supporting this interpretation, social presence increased transitions to stretch-attend and reduced transitions to freezing in dominant mice (Figures 3I–L). Notably, social behaviors were more frequently followed by defensive behaviors under rat exposure than looming (Figures 3Q–T), suggesting that the drive for social support is enhanced under prolonged, multisensory threat. These differences in social modulation of defensive behaviors indicate that social context flexibly adjusts behavioral priorities according to ecological demands: it dampens passive defense and promotes recovery following transient threats while facilitating active coping during sustained predator encounters.

Threat exposure also reshaped social interaction in a rank-dependent manner. Across both paradigms, threat exposure increased social engagement and reinforced social hierarchy, suggesting that enhanced social cohesion represents a common adaptive response to predation risk by reducing individual vulnerability and facilitating collective defense. Notably, social behaviors increased after escape from looming stimuli and withdrawal from the rat, particularly when these actions were initiated by dominant mice, suggesting that dominant individuals serve as key drivers of social recruitment under threat. However, the nature of the two threats likely gives rise to distinct mechanisms of social regulation. A looming stimulus represents an acute, rapidly resolved threat, allowing social interactions to occur primarily after threat exposure. Consistent with this interpretation, looming exposure selectively increased the duration of social behavior in subordinate mice, while dominant mice became more proactive in initiating social interactions. Together with the increased transitions between rearing and freezing, these findings suggest that heightened vigilance promotes rapid post-threat social engagement. In contrast, rat exposure represents a sustained multisensory predator threat that maintains a persistently elevated defensive state, increasing the value of preserving social cohesion during ongoing danger. Accordingly, both the frequency and duration of social interactions increased in dominant and subordinate mice, huddling emerged as the predominant social behavior, and transitions between defensive and social behaviors became more frequent. These findings suggest that social behaviors are tightly integrated with ongoing defensive responses, potentially serving as a strategy for safety seeking and coordinated coping under sustained threat. Thus, looming and rat exposure reveal complementary regimes of social regulation: rapid post-threat social engagement following transient danger and coordinated social coping during sustained predator threat.

Such rank- and threat-dependent modulation is critical for group survival. Under transient threat, stronger fear reduction in dominants may allow leaders to make correct decisions for the group under pressure. Under sustained threat, the shift of dominants toward active defense mirrors natural patterns in which leaders take on greater risk-assessment responsibilities (Blanchard and Blanchard, 1989; Davis et al., 2009). Such role differentiation could enhance group survival by distributing vigilance and risk-taking across the hierarchy. Furthermore, the increase in social interactions following threat exposure likely reflects an evolutionarily conserved coping mechanism that enhances collective security—a phenomenon observed across multiple species, including humans (Morris et al., 1976; Preston et al., 2021; Taylor, 1981; von Dawans et al., 2012).

What neural circuits mediate the social modulation of innate defensive behavior? We propose that the medial prefrontal cortex (mPFC) serves as a central hub in this process. On one hand, mPFC plays a key role in fear regulation (Marek et al., 2013; Sotres-Bayon and Quirk, 2010), social behavior related to dominance hierarchy (Kingsbury et al., 2019; Wang et al., 2011), and is both necessary and sufficient for social buffering of fear responses (Gutzeit et al., 2020; Lungwitz et al., 2014). On the other hand, innate defensive responses to looming and predator threats are mediated by partially distinct subcortical circuits (Silva et al., 2016a), including the superior colliculus (SC) (Evans et al., 2018; Shang et al., 2015; Wei et al., 2015) and the ventromedial hypothalamus (VMHdm) (Kennedy et al., 2020; Silva et al., 2016b), which receive top-down input from the mPFC and are interconnected (Benavidez et al., 2021; Oh et al., 2014; Risold et al., 1997). Therefore, we hypothesize that social context and dominance-related signals are integrated within the mPFC and subsequently modulate activity in defensive circuits to shape threat-specific behavioral strategies.

Conversely, threat may reshape social behavior through partially distinct neural circuits that converge on the mPFC. SC-centered visual threat circuits may facilitate rapid post-threat social engagement by recruiting vigilance-related circuits (Evans et al., 2018; Shang et al., 2018; Wei et al., 2015), whereas VMHdm-centered predator-defense circuits may promote social cohesion by recruiting circuits that support coordinated coping during persistent defensive states (Kunwar et al., 2015; Silva et al., 2016b; Wang et al., 2015). Testing these hypotheses will require future studies that combine circuit-level recordings and causal perturbations within the behavioral framework established here.

One limitation of our study is the exclusive use of male mice. Because females show distinct defensive and social patterns under threat (Blanchard et al., 1991; Pentkowski et al., 2018), extending this paradigm to females with careful control for estrous cycles will provide a more comprehensive understanding of hierarchy-dependent social modulation.

In conclusion, our findings reveal that social presence does not simply suppress fear but selectively tunes defensive and social behaviors according to both rank and threat type. This adaptive flexibility allows groups to manage risk while maintaining cohesion, paralleling collective coping strategies observed in humans and other animals. By providing a behavioral framework for probing the neural basis of social modulation of innate fear, our work opens new avenues for dissecting the interplay between social and defensive circuits. Given that maladaptive fear and social dysfunction often co-occur in neuropsychiatric disorders, these insights may inform mechanistic models of social-affective dysfunction and inspire new translational approaches.

## Materials and methods

### Animal

Male C57BL/6J mice (*n* = 64, 8–12 weeks old) were pair-housed under a 12-h light/12-h dark cycle. Male Long-Evans rats (6 months old) were housed under the same light-dark cycle. All behavioral experiments were performed during the light phase. All experimental procedures complied with animal welfare guidelines and were approved by the Institutional Animal Care and Use Committee at the Chinese Institute for Brain Research, Beijing.

### Arena design

The looming-exposure arena (48 (L) *×* 48 (W) *×* 25 (H) cm) was constructed from infrared-transparent acrylic, and visual stimuli were presented with an overhead monitor (53 *×* 30 cm). A prism-shaped shelter (20 *×* 15 *×* 12 cm) was placed in one corner with the opening facing the monitor. Illumination was provided by two diagonally positioned infrared lamps. For real-time tracking, an OpenMV camera was placed 80 cm below the arena. A second infrared camera (LBAS-U350-74M, LUSTER LightTech) was placed 75 cm below the arena to record animal behavior at 30 Hz.

The rat-exposure arena (38.5 *×* 21 *×* 40 cm) consisted of three opaque acrylic walls and one infrared-transparent wall for video recording. The rat holding cage (18 *×* 14 *×* 35 cm) was made of transparent acrylic, with the wall facing the mouse perforated to allow visual, olfactory, and auditory cues. Behavior was recorded using two cameras: one overhead (75 cm above the floor) and one lateral (50 cm from the infrared-transparent wall).

### Tube test

Social rank within each mouse pair was determined using the tube test (Lindzey et al., 1961; Wang et al., 2011). To minimize stress, each pair was co-housed for at least two weeks with a 15-cm tube placed in their home cage to allow voluntary exploration. Before rank assessment, mice were trained to traverse a 30-cm tube over two consecutive days (10 trials per day), with entry alternating between ends. On the test day, each pair underwent up to seven competitive trials in the same 30-cm tube. A “win” was defined as one mouse advancing through the tube while the opponent retreated completely (all four paws outside) for at least 5 s. The first mouse to achieve four wins was designated as dominant. Social rank stability was reassessed one day after threat exposure; only pairs with consistent ranks were included in subsequent analyses.

### Threat exposure

Looming exposure was conducted over two consecutive days using a counterbalanced design of single and paired trials. Each mouse underwent one 30-minute session per day, beginning with 10 min of free exploration. Looming stimuli were presented on a monitor 25 cm above the arena. Each stimulus consisted of a dark disk on a grey background that expanded from 0*^◦^* to 20*^◦^* of visual angle at 40*^◦^*/s and remained at 20° for 0.25 s. Stimuli were triggered when a mouse entered a circular zone (13 cm in diameter) beneath the monitor center. Each mouse could trigger up to 10 stimuli per session, with a minimum inter-stimulus interval of 2 minutes to prevent habituation to the looming. A total of 40 mice were used.

Rat exposure followed the same two-day counterbalanced design. Each session began with a 5-minute habituation period, followed by 5 minutes of exposure to a live rat. A total of 24 mice were used.

### Animal tracking and locomotor analysis

In the looming paradigm, seven key points on each mouse were tracked using Deeplabcut (Mathis et al., 2018): nose, left ear, right ear, neck, spine, tail base, and tail middle. The model was trained on 1000 labeled frames. Data were filtered for likelihood *>* 0.9 and instantaneous speed *<* 200 cm/s, with missing values linearly interpolated. In the rat paradigm, mouse coordinates were extracted from recorded videos using EthoVisionXT (Noldus).

Arenas were divided into 2 *×* 2 cm grid cells. For each cell, the animal’s velocity, speed, and occupancy time were calculated and averaged across animals. Behavioral zones were identified using a data-driven approach. In the looming arena, the edge zone width was defined at the elbow point of the “time in edge zone” curve (Figure S1A). In the rat arena, the far zone boundary was set at the elbow point of the “time in far zone” curve (Figure S2A), while the near zone boundary was determined at the stable point of the “near-middle zone transition” curve (Figure S2B). Time allocation and speed were subsequently calculated for each zone.

### Behavioral annotation and categorization

Behavioral annotation was performed using BENTO (Segalin et al., 2021), with all labels being mutually exclusive. We focused primarily on defensive and social behaviors, as well as several neutral solitary behaviors related to anxiety and defensive state, including sniffing, grooming, and rearing. Four behaviors were annotated in both paradigms: freezing (complete stillness for *≥* 0.5 s), sniffing (nose twitching while investigating the ground), grooming (using mouth or paws to clean the fur or skin), and tail rattling (vigorous tail shaking in a crouched or alert posture). Paradigm-specific behaviors for looming exposure included up-stretch (extending the head and neck toward the screen with limbs stationary), rearing (lifting the forelimbs off the ground while facing the monitor), and escape (fleeing away from the looming stimulus). For rat exposure, annotated behaviors include approach (moving toward the rat), investigation (nose contact with the rat cage or facing the cage within 5 cm of it), withdrawal (moving away from the rat), and stretch-attend (extending the body toward the rat while remaining stationary). In paired conditions, five social behaviors were annotated: approaching the partner, sniffing the partner (nose-directed investigation of the partner’s face, body, or tail), grooming the partner, following the partner, and huddling (remaining close to the partner without engaging in other social behaviors). All remaining unannotated behaviors were grouped as other neutral solitary behaviors.

Defensive behaviors in the rat exposure paradigm were grouped into two categories: passive and active defense. Passive defense comprised immobility-based strategies, including freezing and tail rattling. Active defense comprised movement- or posture-dependent strategies, including approach, investigation, withdrawal, and stretch-attend.

### Behavior transitions

Sequences of annotated behaviors were extracted for each animal. A transition was defined when two consecutive behaviors occurred within 1 s; intervals of *≥*1 s were classified as transitions through other neutral solitary behaviors. Transition matrices were computed for each animal and averaged across animals for group analysis. Transition difference matrices were generated by subtracting transition probabilities in the single condition from those in the paired condition (Figure 3), or by subtracting transition probabilities during baseline from those during threat exposure (Figure 4), followed by averaging across animals. For visualization, node size was scaled with the total time allocated to each behavior (*T*) according to *r* = 0.01 +0.05 *×* ln(1+*T*), and edge thickness indicated the transition probability.

### Behavioral analysis

To quantify the temporal correlation between dominant and subordinate behaviors, we performed a chi-squared test of independence on the contingency table of observed behavior frequencies. Expected probabilities were calculated under the null hypothesis that the two behaviors are independent by multiplying the respective marginal proportions of dominant and subordinate mice.

To quantify how freezing and investigation were influenced by the behavior of social partners, we aligned freezing or investigation bouts to the onset or offset of specific partner behaviors and quantified the difference within 1s before and after each behavioral event. Statistical significance was assessed using a permutation test, in which the temporal relationship between the focal mouse’s behavior and the partner’s behavior was randomly shuffled.

To quantify social interactions, individual social behaviors occurring within *≤* 1 s were first merged into social episodes. An interaction was defined as a pair of social episodes—one from each mouse—that either temporally overlapped or were separated by a gap of *≤* 1 s. Based on the temporal order of behavior initiation, interactions were classified into six types reflecting asynchronous initiation, with the initiating mouse designated as the proactive individual (Figure 4H). The proactive bout response rate was calculated as the proportion of proactive bouts that received a response.

### Quantification and statistical analysis

No statistical methods were used to predetermine sample size. Data normality was assessed using the Shapiro–Wilk test. For normally distributed data, a two-sample *t*-test or paired *t*-test was applied; otherwise, the Mann–Whitney *U* test or Wilcoxon signed-rank test was applied. For multi-group comparisons, one-way or two-way analysis of variance (ANOVA) with appropriate *post-hoc* tests was used to assess the effects of time and social variables. The Kolmogorov–Smirnov (KS) test was used to compare distributions. Categorical data were analyzed using the chi-squared test or Fisher’s exact test. Permutation tests were used to assess the significance of event-aligned behavioral analyses by comparing the observed statistics with null distributions generated by randomly shuffling temporal relationships between behavioral events. Details of the statistical analyses are provided in the figure legends and the Results section.

## Acknowledgments

Ling-yun Li is supported by the Natural Science Foundation of Beijing Municipality (5244028), the National Natural Science Foundation of China (32471071), and the R&D Program of Beijing Municipal Education Commission (1240030201). Ya-tang Li is supported by the National Natural Science Foundation of China (32271060), the Natural Science Foundation of Beijing Municipality (IS23073), and the start-up fund from CIBR.

## Author contributions

Ling-yun Li and Ya-tang Li supervised the project; Ling-yun Li, Xinjian Gao, and Ya-tang Li designed the experiments; Xinjian Gao collected the data; Xinjian Gao, Ling-yun Li, Jun Zhang, and Wen-wei Wu analyzed the data; Ling-yun Li and Ya-tang Li wrote the manuscript.

## Data and code availability

Data and code are available in a public GitHub repository (https://github.com/YatangLiLab/ Gao-2026-rank-threat-soical-modulation).

## Declaration of interests

The authors declare no competing interests.

## Supplemental information

Additional details related to the Results and Methods sections are provided below.

## Video Titles and Legends

**Videos 1–4:** Example videos showing the four types of behavioral decision in response to looming stimuli: escape, escape after assessment, freezing, and no response.

**Video 5–8:** Example videos showing active and passive defense during live rat exposure.

---

**Figure S1:**
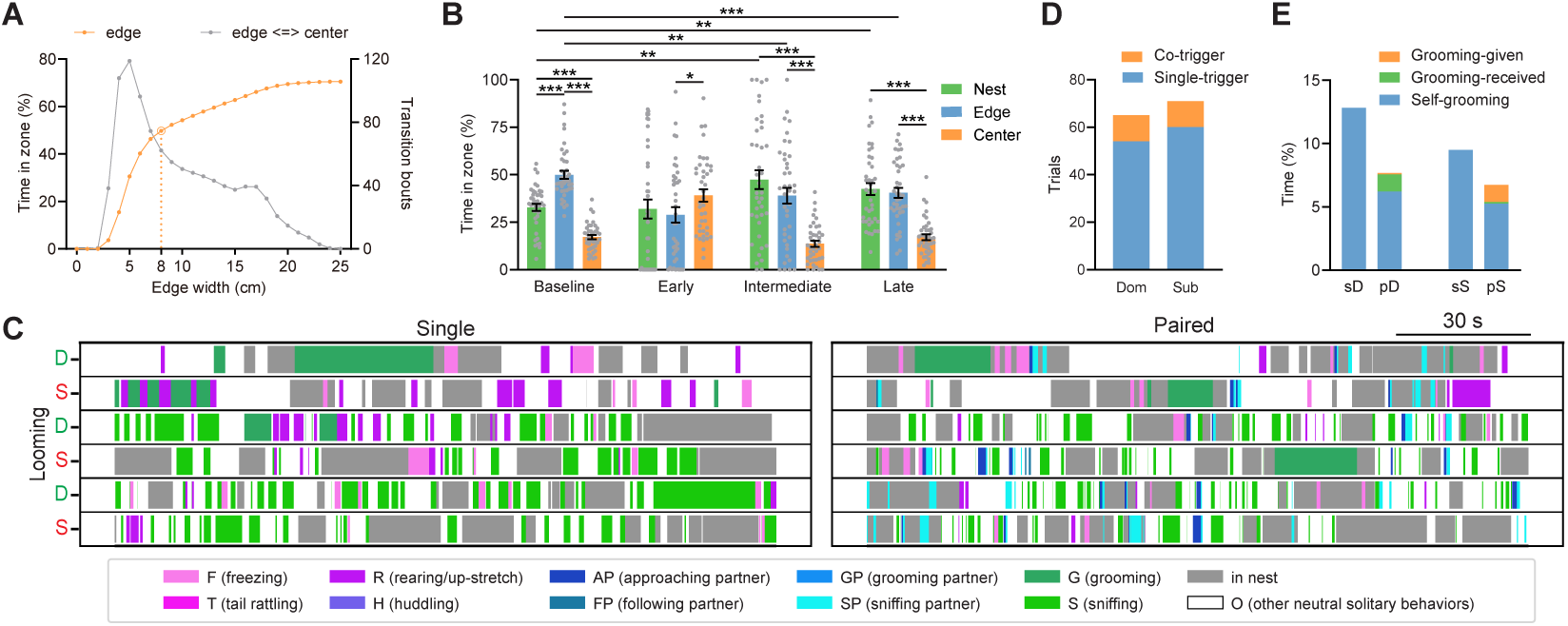
Characterization of looming-evoked defensive behaviors. (A) Time in the edge zone (orange) and transitions between edge and center zones (gray) plotted against edge width. The orange circle marks the elbow point, which was selected as the edge width for analysis in Figure 1. *n* = 40 mice. (B) Time spent in each zone across different time windows under single exposure. *n* = 40 mice; paired *t*-test. (C) Representative raster plots of behaviors during the habituation phase in single and paired looming exposures. (D) Distribution of co-triggered and single-triggered trials across ranks. *n* = 65 bouts (Dom) and 71 bouts (Sub). (E) Time allocation for three types of grooming. *n* = 20 mice for all groups. \**p* < 0.05; ** *p* < 0.01; *** *p* < 0.001.

**Figure S2:**
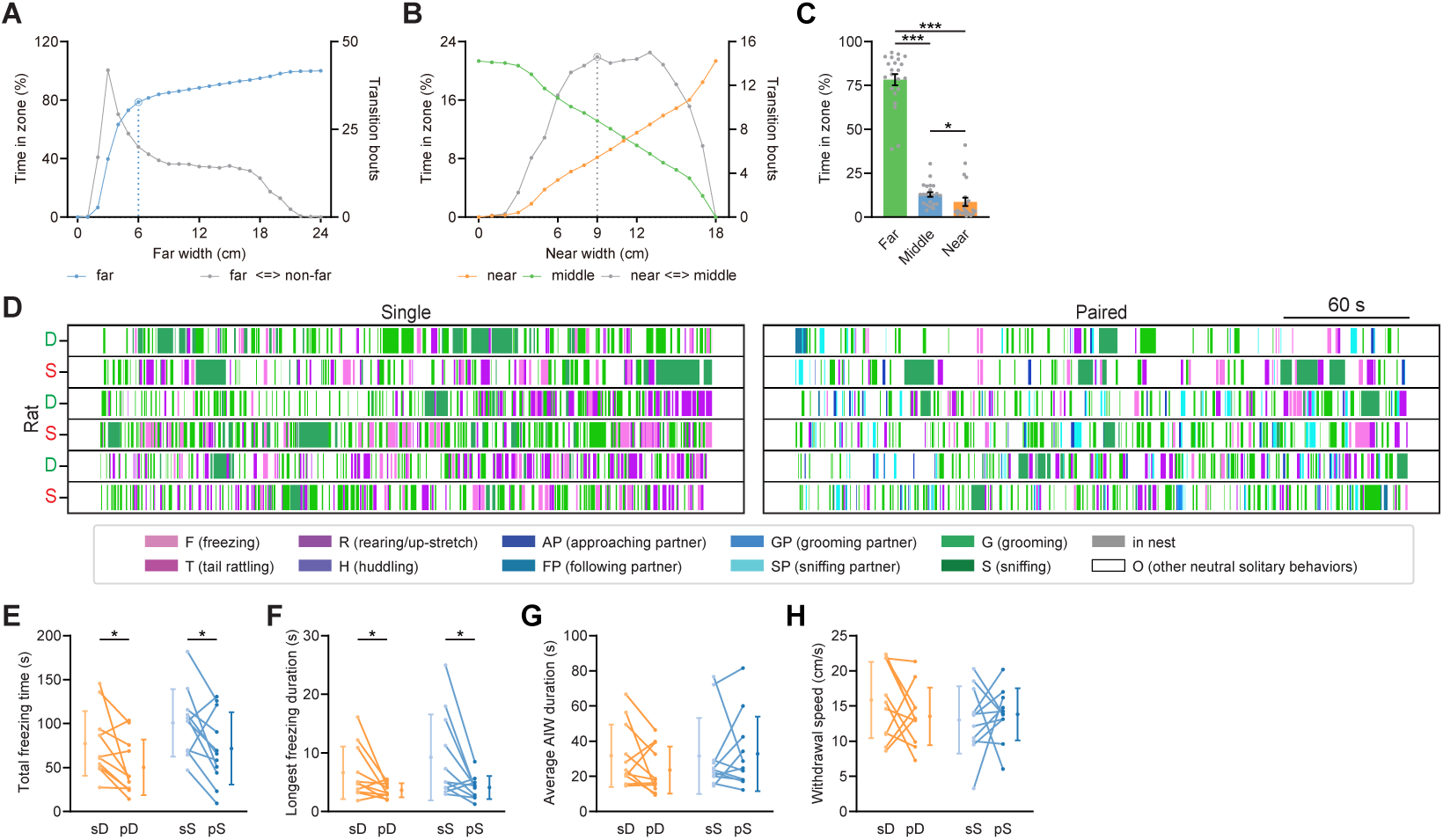
Characterization of rat-evoked defensive behaviors. (A) Time in the far zone (blue) and transitions between the far and the non-far zones (gray) plotted against far zone width. The blue circle marks the elbow point, which was selected as the far zone width for analysis in Figure 2. *n* = 22 mice. (B) Time in the near (orange) and middle (green) zones and transitions between near and middle zones (gray) plotted against near zone width. The gray circle marks the stable point, which was selected as the near zone width for analysis in Figure 2. *n* = 22 mice. (C) Percentage of time spent in each zone during rat exposure. Paired *t*-test; *n* = 22 mice. (D) Representative raster plots of behaviors during the habituation phase in single and paired rat exposures. (E) Total freezing time. Paired *t*-test; *n* = 11 mice for each group. (F) Longest freezing duration. Paired *t*-test; *n* = 11 mice for each group. (G) Average AIW duration. *n* = 11 mice for each group. (H) Withdrawal speed. *n* = 11 mice for each group. \**p* < 0.05; ** *p* < 0.01; *** *p* < 0.001.

**Figure S3:**
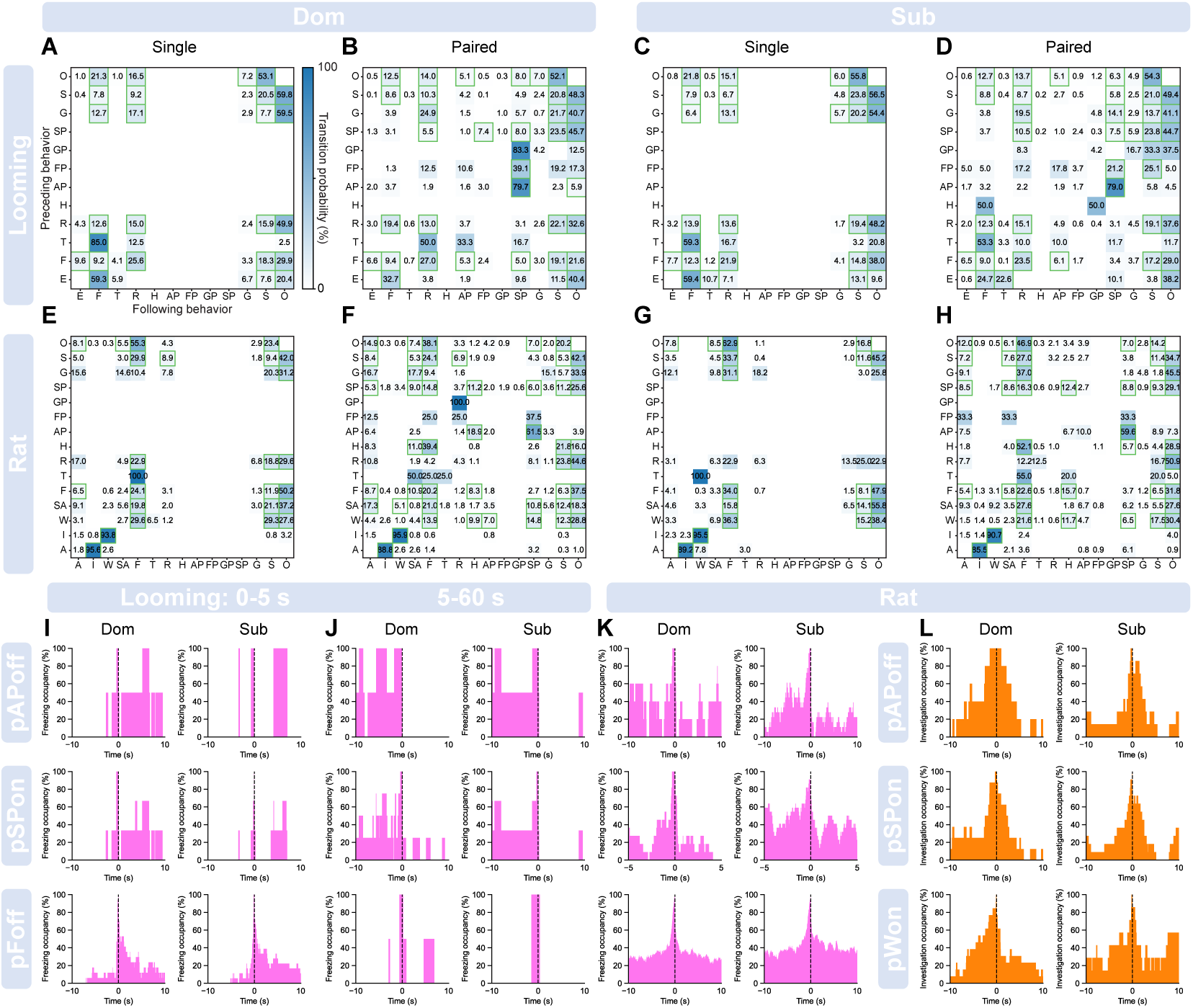
Transition probabilities between behavior pairs and behavior occupancy aligned to partner’s associated behaviors. (A–B) Transition probability matrices for single (A) and paired (B) exposure in the looming paradigm for dominant mice. *n* = 20 mice. Green squares indicate transitions observed in more than five mice. (C–D) Same plot as A–B for subordinate mice. *n* = 20 mice. (E–H) Same plot as A–D for rat exposure paradigm. *n* = 11 mice for both dominant and subordinate groups. (I–J) Freezing occupancy aligned to partner’s associated behavioral events (pAPoff, top; pSPon, middle; pFoff, bottom) for dominant and subordinate mice during different time periods in looming exposure (I: 0–5 s post-looming; J: 5–60 s post-looming). *n* = 20 mice. (K) Same plot as I–J for freezing occupancy in rat exposure. *n* = 11 mice. (L) Investigation occupancy aligned to partner’s associated behavioral events (pAPoff, top; pSPon, middle; pWon, bottom) for dominant and subordinate mice during rat exposure. *n* = 11 mice.

**Figure S4:**
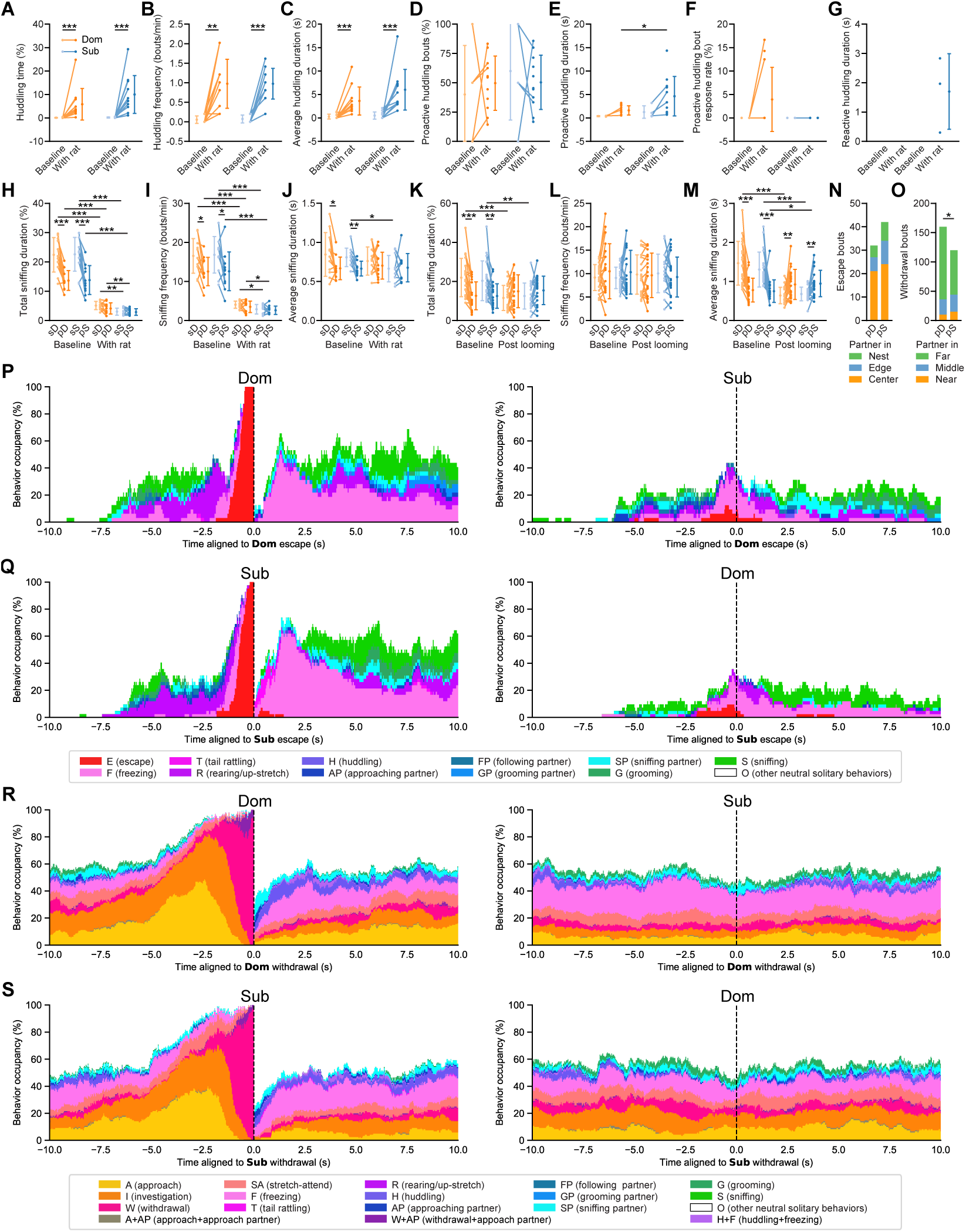
Threat exposure reshapes social interactions and dynamics. (A–C) Total huddling time (A), huddling frequency (B), and average huddling duration (C) during the 5-min periods before and during rat exposure. *n* = 11 mice for both groups, paired *t*-test. (D-G) Percentage of proactive huddling bouts (D), proactive huddling duration (E), response rate (F), and reactive huddling duration (G) during the 5-min periods before and during rat exposure. *n* = 11 mice for both groups, paired *t*-test. (H–J) Total sniffing duration (H), sniffing frequency (I), and average sniffing duration (J) during the 5-min periods before and during rat exposure. *n* = 11 mice for both groups, paired *t*-test. (K–M) Same plot as H–J during a 1-min period before and after looming exposure. *n* = 22 mice for both groups, paired *t*-test. (N) Escape bouts categorized by partner’s location at escape offset during paired looming exposure. *n* = 32 bouts (Dom) and 42 bouts (Sub). (O) Withdrawal bouts categorized by partner’s location at withdrawal offset in paired rat exposure. *n* = 160 bouts (Dom) and 120 bouts (Sub), chi-squared test. (P) Behavior occupancy aligned to dominant escape onset in paired looming exposure. Left, dominant mouse; right, subordinate mouse. *n* = 32 bouts. (Q) Behavior occupancy aligned to subordinate escape onset in paired looming exposure. Left, subordinate mouse; right, dominant mouse. *n* = 42 bouts. (R) Behavior occupancy aligned to dominant withdrawal onset in paired rat exposure. Left, dominant mouse; right, subordinate mouse. *n* = 160 bouts. (S) Behavior occupancy aligned to subordinate withdrawal onset in paired rat exposure. Left, subordinate mouse; right, dominant mouse. *n* = 120 bouts.

## Notes

### Competing Interest Statement

The authors have declared no competing interest.

### Summary of Updates

We have substantially revised the manuscript and regenerated all figures.

